# Ultralight Solar Transmitter Enables Fine-Scale Movement Ecology in North American Hummingbird Migration

**DOI:** 10.64898/2025.12.29.696924

**Authors:** Emma M. Rhodes, Sean Burcher, Eric Johnson, Sarahy Contreras-Martínez, David A. La Puma, Kyle Shepard, DeAnna Williams, Michael Lanzone

## Abstract

Due to attachment weight limitations, animals under 5g have remained a major methodological barrier in movement ecology, limiting tracking options for many small-bodied organisms.
We developed and field-tested an ultra-light solar powered transmitter (BlūMorpho 0.058-0.060g) designed to enable continuous, near real-time tracking using a broad, crowd sourced network.
We validated the system on two North American hummingbird species during spring migration, demonstrating reliable detection, multi-week retention, broad spatial coverage (United States, Mexico and Canada), and long-distance migration paths including a 6,526km cumulative track.
Our results highlight the ability to track the smallest of species across the full migratory range with fine-scale spatial resolution and high temporal resolution using the BlūMorpho transmitter. Importantly, we present the first migratory tracks known to exist on North American hummingbird species.

**Data/Code for peer review statements:** R Code used for all analyses is attached for peer review.

## 1 INTRODUCTION

Migration is a central ecological phenomenon, yet its transient and large-scale nature makes it inherently difficult to study. Every year, billions of birds undergo annual migratory movements driving questions that span conservation (Fraser et al., 2018), ecology (Rousseau et al., 2020), evolution (Gu et al., 2024), speciation (Williamson et al., 2024), and physiology (Rhodes et al., 2024). The study of migration in recent decades has created its own subdiscipline of movement ecology, which can be defined as the study of when, where, why, and how organisms move (Nathan et al., 2008). Technological advancements have transformed the field, providing insight into behaviors previously undocumented.

Numerous studies in avian movement ecology have documented diverse life histories and migratory strategies. Yet a major gap remains; tracking the smallest animals (Wikelski et al., 2007). Device weight has been the key limiting factor, preventing the application of high-resolution tracking technologies on species like North American hummingbirds that can weigh less than 10g (McKinnon & Love, 2018). To date, the smallest telemetry devices have been primarily data loggers, including geolocators and archival GPS tags, which require physical recovery to access stored data and may have spatial error (ranging from 100-1000 km) (McKinnon & Love, 2018). Existing VHF and UHF transmitters do not require retrieval but require maintaining stationary receivers on a large-scale (Taylor et al., 2017). This makes them unsuitable for studying fine-scale, long-distance movements for species which recapture is impractical or impossible. Real-time transmitters capable of remote data delivery have traditionally required bulky batteries, limiting their use to larger-bodied animals (Wikelski et al., 2007). The BlūMorpho (Cellular Tracking Technologies (CTT), Rio Grande, NJ, USA) transmitter breaks this trade-off by combining solar power with near real-time transmission via a ubiquitous crowd-sourced network; all in a device weighing just 0.058–0.070 g. This enables continuous, remote tracking at high spatial and temporal resolution, with no need for recovery. Here, we report the application of a solar-powered transmitter (BlūMorpho, deployed on two hummingbird species, Rufous (*Selasphorus rufus*) and Ruby-throated (*Archilochus colubris*), during spring migration along the Northern Gulf of Mexico (Bassett & Dietrich, 2021; Weidensaul et al., 2020).

## 2 MATERIALS AND METHODS

### 2.1 Data Collection

Fieldwork was conducted in March and April of 2025 during the initiation of spring migration along the Northern Gulf of Mexico, including Coastal Alabama and the Florida Panhandle (U.S.). Hummingbirds were selected for transmitter deployment given 1.) their small size and lack of available migration data and 2.) because they are known to migrate long-distances though fine-scale migration movement documenting these migrations are limited. Ruby-throated Hummingbirds (*Archilochus colubris)* and Rufous Hummingbirds (*Selasphorus rufus*) were selected since they can be found throughout the Gulf Coast during our trapping period. We also utilized already established hummingbird banding sites previously reported (Bassett & Cubie, 2009; Bassett & Dietrich, 2021). Both species were targeted for transmitter deployment using two Bassett traps following standard protocols (Bassett & Cubie, 2009). Capture efforts occurred opportunistically, between sunrise and sunset. Individuals were captured, banded, and telemetered with all appropriate permits: USGS Bird Banding Permit (#24217), Alabama (#2025104370668680 & #2025104369668680), and Florida (#LSSC-22-00033). Birds were aged based on bill corrugations and sexed by plumage (Ortiz-Crespo, 1972; Pyle, 1997). Fat and muscle scores (0-3) were assigned following previously reported methods (Rhodes et al., 2024). Wing chord and exposed culmen were measured with digital calipers and tail length with a ruler in mm. Birds were weighed using a digital scale and weights were recorded to a tenth of a gram. Transmitters were deployed only on individuals for which the weight of all attachments did not exceed 3% of body mass following standard protocol (McKinnon & Love, 2018). We initially aimed to tag both species at each site; however, this was achieved at only three of the seven sites due to species availability.

### 2.2 Attachment methods

We trimmed a ∼3x10mm area of feathers below the nape and applied a small amount of either cyanoacrylate adhesive (Loctite Super Glue Gel Control) or eyelash adhesive (Velour Strong Eyelash Glue Brand) to the back side of the transmitter and gently pressed into place (Figure 1). Once adhered and dry, transmitter function was confirmed using a handheld receiver (Cellular Tracking Technologies, Rio Grande, NJ). Birds were released at the site of capture. Details on placement and attachment is provided in a step-by-step format in the Supporting Information (See Figure S1).

**FIGURE 1.**
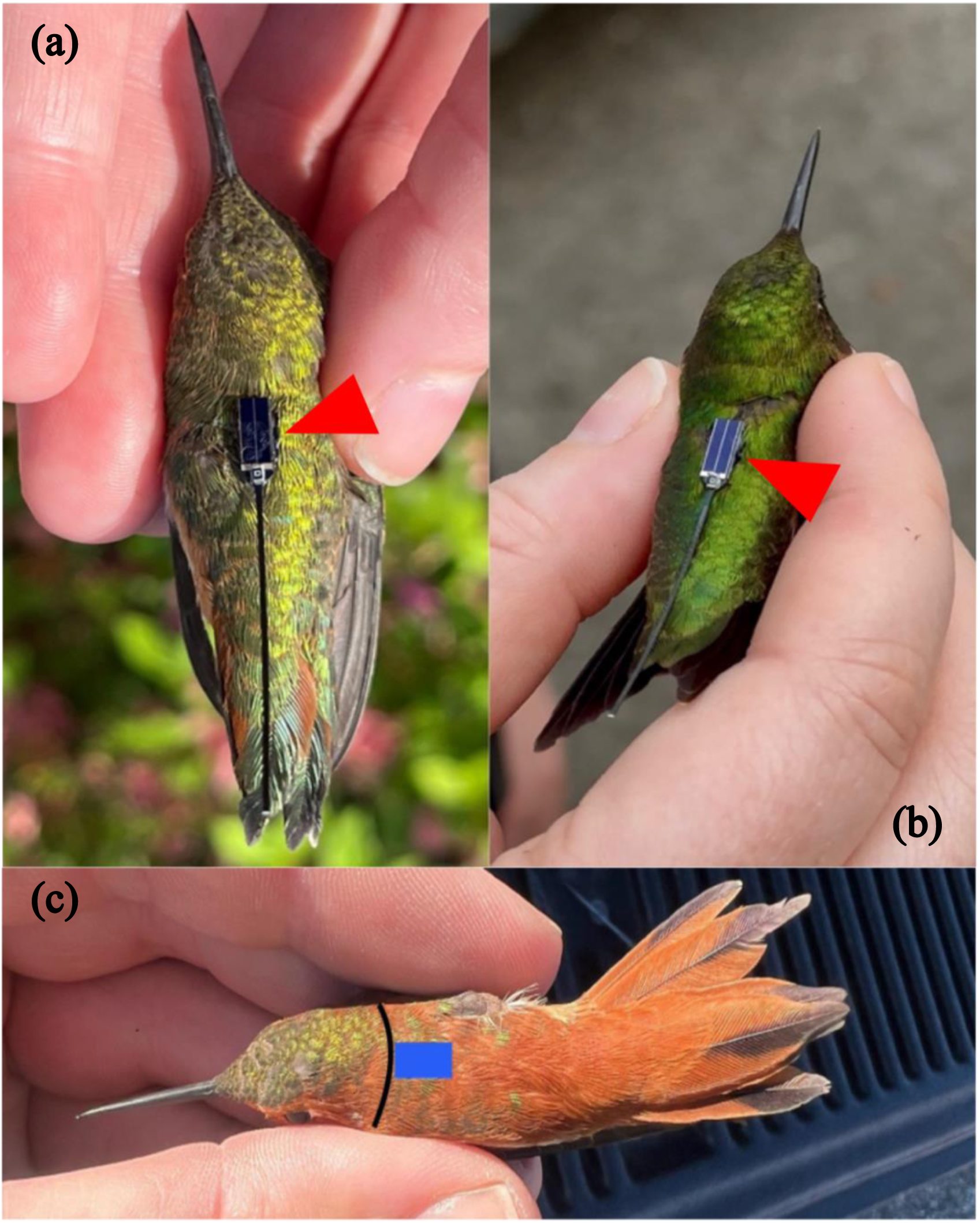
Images of a telemetered Rufous Hummingbird (*Selasphorus rufus*) (a) and Ruby-throated Hummingbird (*Archilochus colubris)* (b) illustrating the attachment location on the center, dorsal side of the animal with a red arrow pointing to the placement of the tag. Panel (c) shows the target area to be trimmed indicated by a blue square. The black line on the bird’s nape indicates the upper-boundary for where trimming and tag placement should be avoided since placing tags on or above this line leads to pulling of the upper-nape feathers creating tension and allows birds to easily pluck and remove feathers leading to shorter tag-retention time.

### 2.3 BlūMorpho transmitter

The BlūMorpho transmitter is digitally coded, solar-powered, and emits a unique 2.4 GHz signal every 3 seconds. Unlike data loggers, it stores no onboard data but leverages a broad network of IoT-enabled receivers as well as Motus-compatible CTT SensorStations (Taylor et al., 2017), Sidekick handheld device, the community-science Terra network (terralistens.com), and most modern smartphones, tablets and computers where both location and Bluetooth services are running in the background. The network polls for new detections every 5 minutes, allowing researchers near real-time access to high-resolution movement data. Because it is solar-powered, tracking is limited to daylight hours.

The data we report are primarily sourced from detections that occurred on existing crowd-sourced location networks. By leveraging the expansive reach of the crowd-sourced location networks, it is possible to collect fine-scale location data across vast regions without the need for the deployment or maintenance of any specialized receiver infrastructure. However, detection data is only collected when the transmitter is within range of a device that is part of an existing crowd-source location network and therefore the data may be biased to areas of higher population densities. All detections were stored in CTT’s user portal from which the data were exported in csv format for analyses. The movement data were analyzed using R (version 4.4.2) (R Core Team, 2023) and mapped in ggplot2 (Wickham, 2016).

## 3 RESULTS

### 3.1 Tag performance and retention

We telemetered 11 individuals across seven sites (Table 1). The average detection duration was 35 days (range: 3 - 121 days) (Table 1). The average distance from release to last detection was 1,032 km ± 1,367 km with a range of 0.03 km to 4730 km. The relatively short retention time is expected, given the temporary adhesive used; environmental exposure and molt likely lead to cases of early detachment. Detection rates immediately following release was 100% and 73% for detections outside the original capture state. Nine of the 11 individuals were detected over 10km from the original capture location (Table 2). Local movement data is biased toward crowd-sourced receivers but produced fine-scale detections from ∼100-300m distance (Figure 2).

**FIGURE 2.**
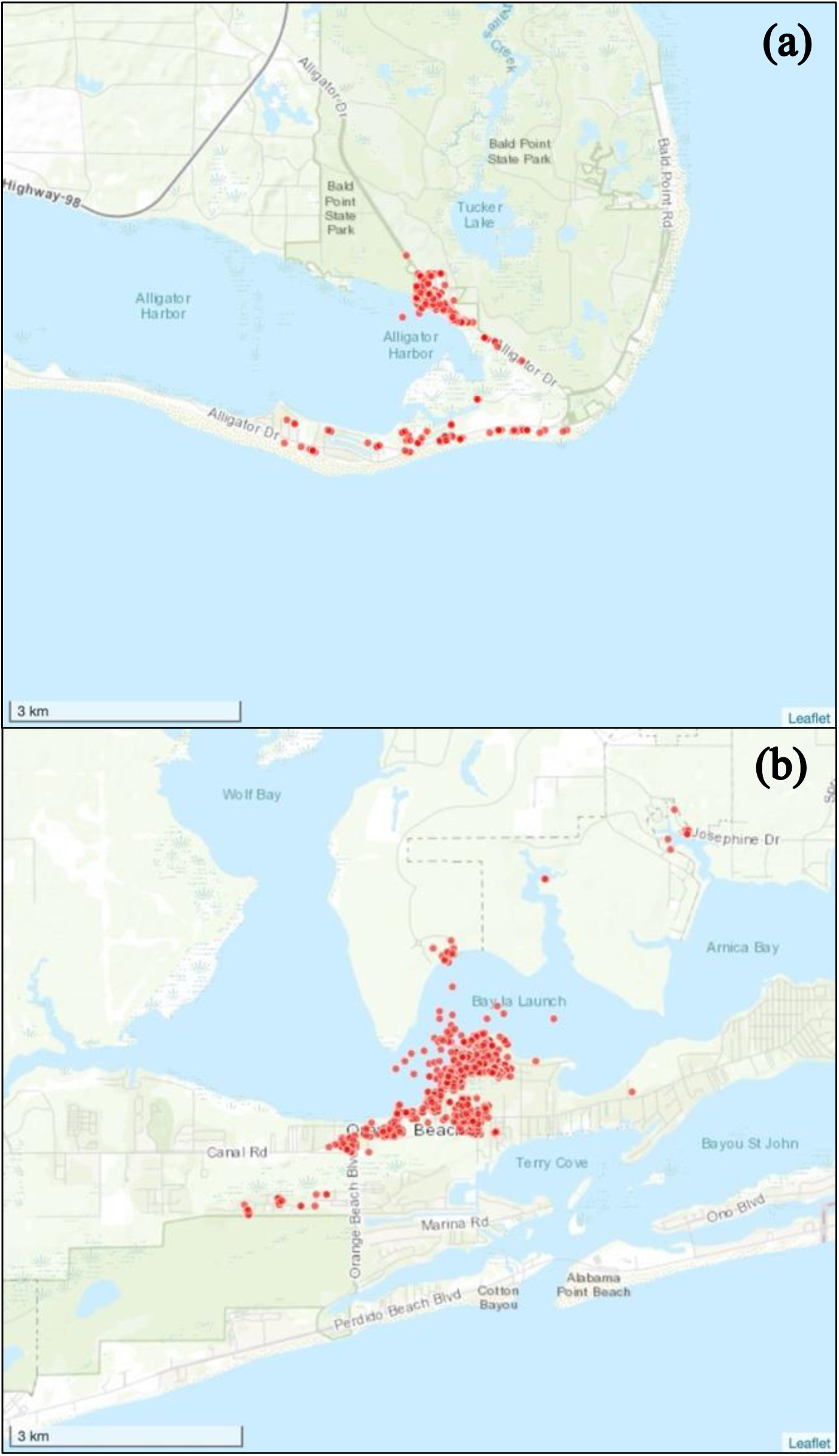
(a) Bird ID F09545 demonstrating local movement in Alligator Point, FL (Franklin County). b) Bird ID F09544 in Orange Beach, AL (Baldwin County). The uncertainty for any location should be aken as the maximum detection range of the digital device that recorded the detection (∼100-300 meters).

**TABLE 1.** Details of 11 individuals of the two species (Rufous and Ruby-throated Hummingbird) with the accompanying transmitter metadata including Bird ID (USGS federal band number), Start and End detections by latitude/longitude, deployment date/times (in coordinated universal time), County/state of capture, date/time of last detection, and location of last detections.

| Bird ID | Start Lat, Long | Date Deployed | Time Deployed (UTC) | County | State | End Lat, Long | Date of Last Detection | Time of Last Detection (UTC) | County | State |
| --- | --- | --- | --- | --- | --- | --- | --- | --- | --- | --- |
| **Rufous Hummingbird (*Selasphorus rufus*)** |  |  |  |  |  |  |  |  |  |  |
| F34924 | 30.479183<br>-87.90676 | 3/27/25 | 11:34 | Baldwin | AL | 34.380088<br>-93.136524 | 4/3/25 | 18:06 | Hot Spring | AR |
| F34925 | 30.515368<br>-87.90494 | 3/27/25 | 13:52 | Baldwin | AL | 30.517574<br>-87.912112 | 3/29/25 | 20:36 | Baldwin | AL |
| F34926 | 30.357192<br>-87.47209 | 3/27/25 | 17:33 | Baldwin | AL | 58.382759<br>-134.736715 | 5/3/25 | 16:08 | Juneau | AK |
| F34928 | 30.515368<br>-87.90494 | 3/30/25 | 20:46 | Baldwin | AL | 38.664631<br>-76.674744 | 4/21/25 | 15:43 | Calvert | MD |
| **Ruby-throated Hummingbird (*Archilochus colubris*)** |  |  |  |  |  |  |  |  |  |  |
| F09544 | 30.479183<br>-87.90676 | 3/27/25 | 11:45 | Baldwin | AL | 30.479426<br>-87.906464 | 4/22/25 | 20:18 | Baldwin | AL |
| F34927 | 30.357192<br>-87.47209 | 3/27/25 | 18:11 | Baldwin | AL | 29.611185<br>-82.529247 | 5/21/25 | 13:05 | Alachua | FL |
| F09545 | 29.912242<br>-84.36708 | 4/11/25 | 15:54 | Franklin | FL | 29.912156<br>-84.367374 | 8/16/25 | 11:54 | Franklin | FL |
| F09546 | 30.502517<br>-86.46218 | 4/11/25 | 23:29 | Okaloosa | FL | 39.576080<br>-82.206508 | 4/30/25 | 23:41 | Perry | OH |
| F09547 | 30.502517<br>-86.46218 | 4/11/25 | 23:47 | Okaloosa | FL | 46.705827<br>-93.194416 | 5/17/25 | 23:06 | Clark Township | MN |
| F34929 | 30.302956<br>-87.56308 | 4/17/25 | 14:16 | Baldwin | AL | 34.502806<br>-80.907859 | 6/9/25 | 18:45 | Fairfield | SC |
| F09548 | 30.302956<br>-87.56308 | 4/19/25 | 13:49 | Baldwin | AL | 29.708374<br>-91.220332 | 8/17/25 | 14:04 | St. Mary Parish | LA |

**TABLE 2.**
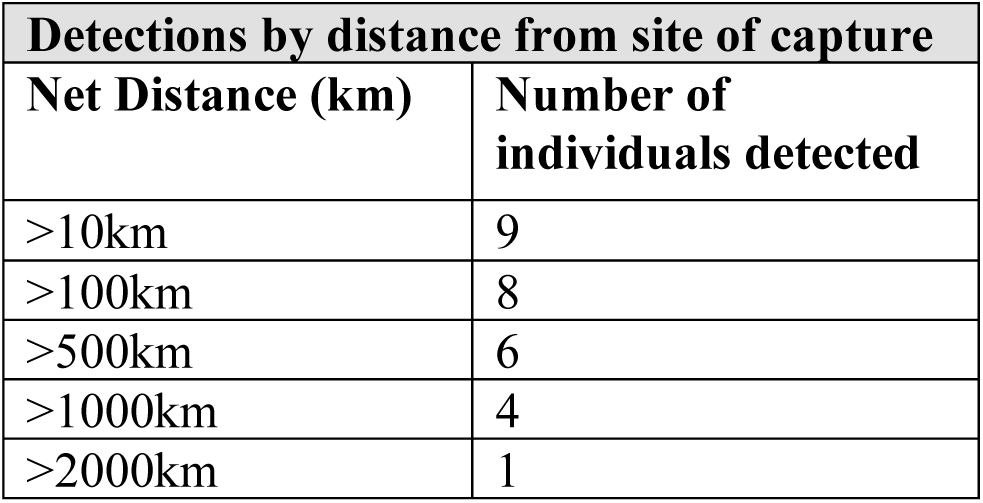
Number of the 11 individuals that were detected over 10, 100, 500, etc. km from original site which they were captured and released.

### 3.2 Summary of migratory routes

Two Rufous Hummingbirds (F34926, F34924) initially exhibited western movements. F34926 continued west, crossing Northern Mexico, then headed north across the Pacific coast to Alaska (Figure 3A). In contrast, we stopped detecting F34924 once it reached the Ozark mountains in Arkansas. Our third Rufous F34928 initially headed northward and then exhibited a serpentine-like track traveling throughout several Eastern U.S. states until it was no longer detected (Figure 3B). Four Ruby-throated Hummingbirds (F09546, F09547, F34927, F34929) exhibited movement aligning with spring migration across the Eastern U.S. (Figure 3B). Three individuals (F09546, F09547, F34929) exhibited direct movements north and northeast towards their breeding grounds (Figure 3B). F09548 stayed near the original capture location until fall migration where it moved westward to Louisiana where it was last detected (Figure 3A). Notably, the Ruby-throated F34927 deviated from the expected north to northeastern spring movement and traveled southeast from its original capture location in Coastal Alabama and moved into Florida, in Alachua County, where it was last detected on 21 May 2025. Two of the seven telemetered Ruby-throated Hummingbirds (F09544, F09545) displayed only local movements surrounding the original capture location thus are not shown in Figure 3 (see Table 1 for details).

**FIGURE 3.**
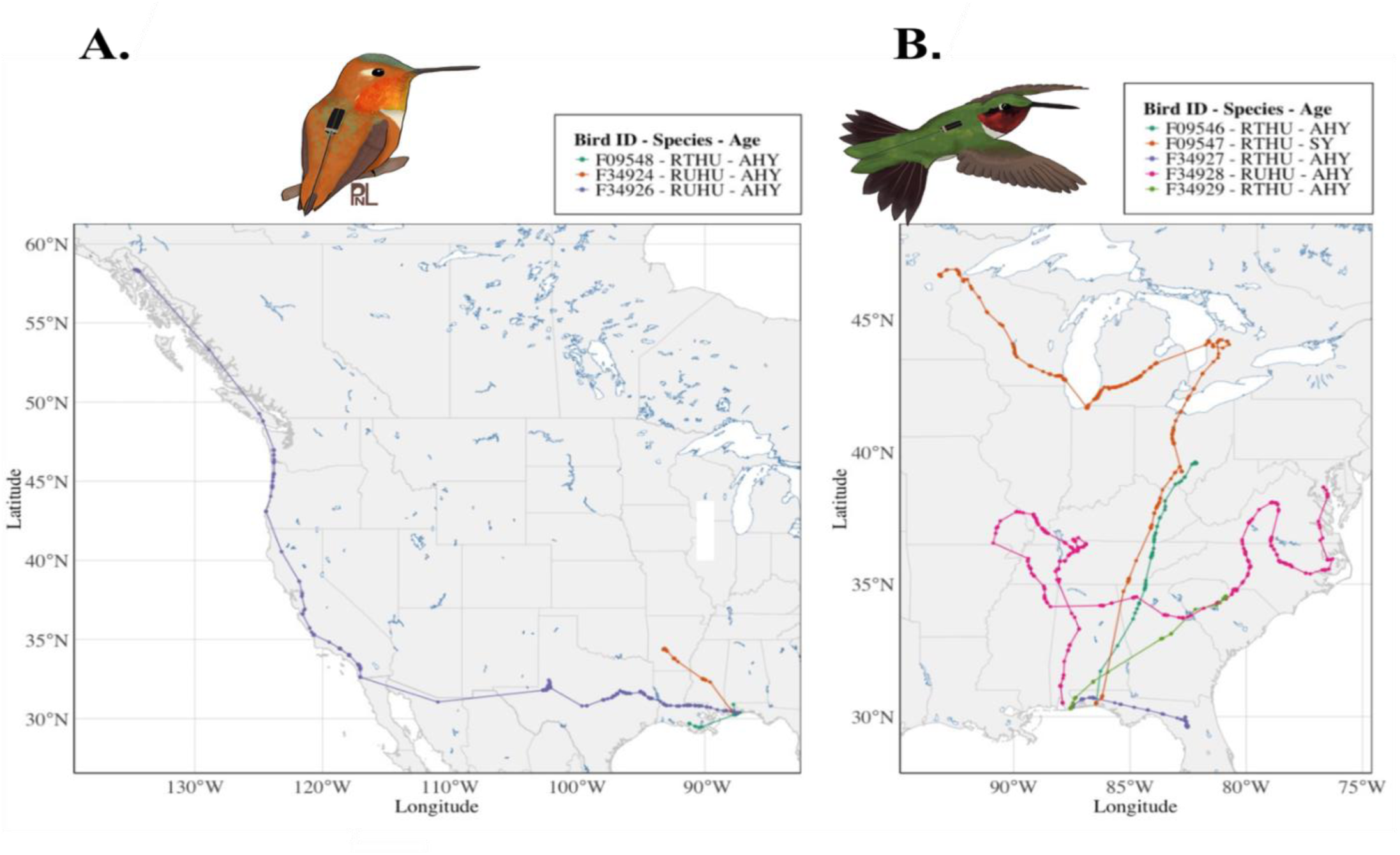
Map of the 11 individual tracks to observe large-scale movements separated out in two panels (A - B.) Bird ID is represented as the band number and species is abbreviated as follows: Ruby-throated Hummingbird (RTHU) and Rufous Hummingbird (RUHU). Age is also given as either AHY (After-hatching-year) or SY (Second-year). X-axis is longitude and Y-axis is latitude by degrees

The longest recorded migration was from a Rufous Hummingbird (F34926) released in Lillian, AL on 27 March 2025 and last detected on 3 May 2025 in Juneau, AK (Figure 3A). The total cumulative migration distance, calculated as the sum of all segments between consecutive detections, was 6,526 km, with a straight-line distance of 4,730 km (Figure 3A). In total, from our 11 individuals, detections occurred across 24 U.S. states, northern Mexico (Sonora) and parts of Canada (British Colombia).

### 3.3 Comparison to previous and current methodology

Traditional banding data from the U.S. collected between 1960-2024 produced a 0.14% recapture rate for Rufous (209 recaptured individuals/151,772 total individuals banded) and 0.03% recapture rate for Ruby-throated Hummingbird (151/501,916) (Database: *USGS Bird Banding Laboratory*, 2024). For the purposes of directly comparing to our dataset, we used Ruby-throated and Rufous recapture data reported by the Bassett and Dietrich (2021) study. We first made a distinction between local recaptures, which we defined as individual birds re-encountered by Bassett and Dietrich (2021). We then defined foreign recaptures as any recapture of individuals banded by Bassett and Dietrich (2021) but was re-encountered from another permitted bird bander. This resulted in 573 local recaptures and 21 foreign recaptures in the Bassett and Dietrich (2021) dataset containing data spanning from 1999 and 2020 (Figure 4). For mapping purposes, all the local recapture data where individuals were detected less than a 100km from the original banding site were filtered out.

**FIGURE 4.**
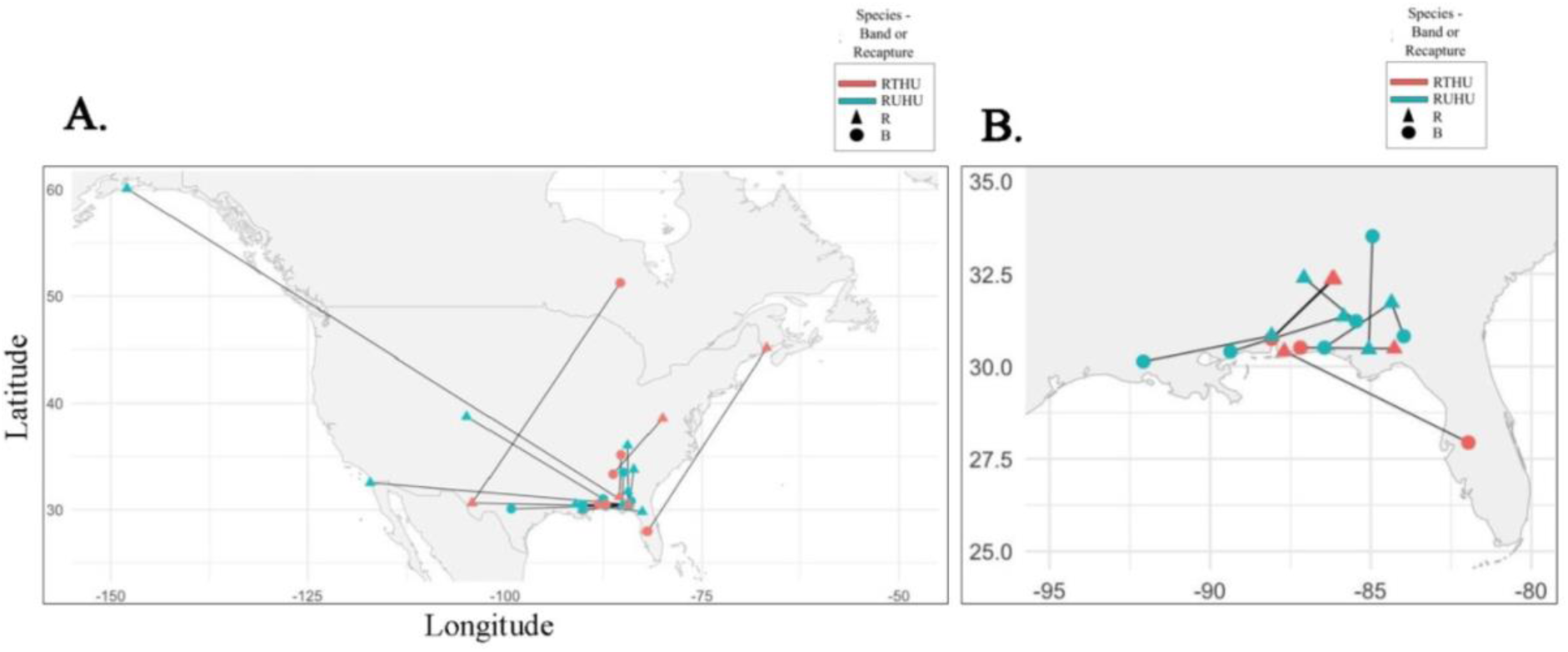
Map of foreign recaptures (a) and local recaptures (b) reported by Bassett and Dietrich (2021). Circles represent the original banding location and triangles, the subsequent recapture locations. Ruby-throated Hummingbird (RTHU) recaptures are represented as pink and Rufous Hummingbird (RUHU) as blue. X-axis is longitude and Y-axis is latitude by degrees

## 4 DISCUSSION

The primary goal of this work is to demonstrate how the application of this cutting-edge tracking technology, leveraging ubiquitous Bluetooth-enabled device networks, will allow researchers to collect vital tracking data on the smallest of animals, such as hummingbirds, in a historically unprecedented level of detail. The ability to track the local and long-distance movement of small species at a fine spatial scale will allow scientists and conservationists to address questions about species’ life cycle stages that were previously inaccessible. Furthermore, this technology has broader applications to track a plethora of small organisms such as the migratory Monarch Butterflies (*Danaus plexippus*). It should be noted that this methodology works exceptional well for diurnal movements but is innately limited to day-time detections since it solar-powered and has no on-board battery. It should also be noted that detections are biased towards areas with IoT digital devices, thus reception of these transmitters is positively correlated with increasingly populated areas.

In this study, we applied this novel technology to two species of hummingbirds where all telemetered individuals weighed less than 5g. Previous hummingbird work utilizing traditional mark-recapture methods such as banding (Rousseau et al., 2020), RFID tags (Zenzal et al., 2014), and stable isotopes (Koehler et al., 2025; Moran et al., 2013), has provided critical insight and baseline information. Yet, they do not provide high enough data resolution to understand individual and population-level variation in migration patterns. While many individuals have been re-encountered at original banding locations, historical banding records have produced very limited foreign recapture data, limiting our ability to fully document hummingbirds’ full annual cycle. For example, Bassett and Dietrich (2021) reported only 1 Rufous Hummingbird foreign recapture following a 20-year period. This individual was originally banded in Florida on its winter grounds and subsequently captured on its presumed breeding grounds in Alaska (Bassett & Dietrich, 2021). Our results highlight the improvement of data collection on hummingbirds through use of a novel transmitter rather than traditional banding alone given 100% of individuals were detected post-release and some individuals were detected hundreds to thousands of km from capture location.

Previous RFID tags deployed on Ruby-throated Hummingbirds along the Alabama coast have provided local feeder information such as feeding rates and possible time of southbound coastal departure but was limited to only presence/absence at feeder traps (Zenzal et al., 2014). In contrast, we demonstrate the ability to pinpoint locations of individuals within a ∼100-300m range with these transmitters.

Previous modeling techniques have also given insight into these species’ migratory behavior (Rousseau et al., 2020); however, these estimates will be improved with the incorporation of individual migration tracks such as the ones presented in our results (Buderman et al., 2025). With both Rufous and Ruby-throated Hummingbirds experiencing recent population declines (English et al., 2021), this new technology will provide critical insights for species management and conservation. Importantly, our results demonstrate the capability of this technology to determine migratory movements, breeding and winter ranges, and stopover ecology. Our continued efforts will allow us to elucidate and further document species-specific patterns, intra-annual migratory strategies, and demographic patterns. Future work will also allow for population-level discoveries such as the extent of migratory connectivity from the breeding and non-breeding grounds of both species.

In conclusion, we demonstrate how a novel lightweight transmitter, the BlūMorpho, unlocks near real-time, fine-scale tracking in hummingbirds—species previously too small for such technology. While this initial effort relied on short-term deployments using glue-on attachment methods, future work should include harnessing techniques which will allow tracking over the full annual cycle. While a harness does exist for larger-bodied hummingbirds (Williamson & Witt, 2021), thorough testing for harnesses is needed for hummingbirds that weigh 5 grams or less.

## Acknowledgments

We thank the University of Guadalajara – CUCSUR and CONANP for supporting the initial fieldwork tests. Thanks to Robert R. and Martha Sargent (the Hummer/Bird Study Group), Fred Bassett and Fred Dietrich (Hummingbird Research, INC), and Kelly Bryan. This work could not have been completed without the special experimental authorization provided by the USGS Bird Banding Laboratory. Special thanks to Antonio Celis-Murillo, Matthew Rogosky, and T.J. Zenzal for providing initial feedback on these methods. Thanks to Wendy Hood, Geoff Hill, and Yufeng Zhang for providing the lead author support. Thank you to Phoebe Lanzone for contributing artwork. Funding for this work was supported by Cellular Tracking Technologies and Banding Coalition of the Americas (BCA).

## Data Availability Statement

Data will be made publicly available at time of publication.

## Author Contributions

E.M.R, S.B., E.J., S.CM., D.A.L., K.S., D.W., and M.L. conceived the ideas and designed the methodology; E.M.R. and K.S. collected the data; E.M.R., S.B., E.J., D.A.P., K.S., and M.L. analyzed the data; E.M.R, K.S., and D.A.L. led the writing of the manuscript. All authors contributed critically to the drafts and gave final approval for publication.

