## Supporting Information for "Ultralight Solar Transmitter Enables Fine-Scale Movement Ecology in North American Hummingbird Migration"

**TABLE 1** Second-year (SY). Fat and Muscle scores were assigned a score 0-3. Mass is in grams and morphometrics are all given in mm.

| Bird ID | Age | Sex | Fat Score | Muscle Score | Mass (g) | Wing (mm) | Tail (mm) | Exposed Culmen (mm) |
|---|---|---|---|---|---|---|---|---|
| **Rufous Hummingbird *(Selasphorus rufus)*** | | | | | | | | |
| F34924 | AHY | M | 1 | 2 | 3.3 | 39.4 | 29 | 16.3 |
| F34925 | AHY | F | 1 | 2 | 3.4 | 41.7 | 26 | 17.8 |
| F34926 | AHY | M | 1 | 2 | 3.8 | 39.8 | 29 | 14.8 |
| F34928 | AHY | M | 3 | 2 | 4.3 | 41.7 | 29 | 17.5 |
| **Ruby-throated Hummingbird *(Archilochus colubris)*** | | | | | | | | |
| F09544 | AHY | M | 0 | 2 | 2.9 | 38.8 | 28 | 15.9 |
| F34927 | AHY | F | 0 | 1 | 3.4 | 43.9 | 27 | 18.7 |
| F09545 | AHY | M | 1 | 2 | 2.7 | 38.6 | 27 | 16.9 |
| F09546 | AHY | M | 0 | 2 | 3.3 | 39.0 | 27 | 16.1 |
| F09547 | SY | M | 2 | 2 | 3.8 | 39.6 | 27 | 15.4 |
| F34929 | AHY | F | 1 | 1 | 3.2 | 43.7 | 26 | 19.6 |
| F09548 | AHY | M | 0 | 1 | 2.8 | 39.3 | 27 | 16.8 |

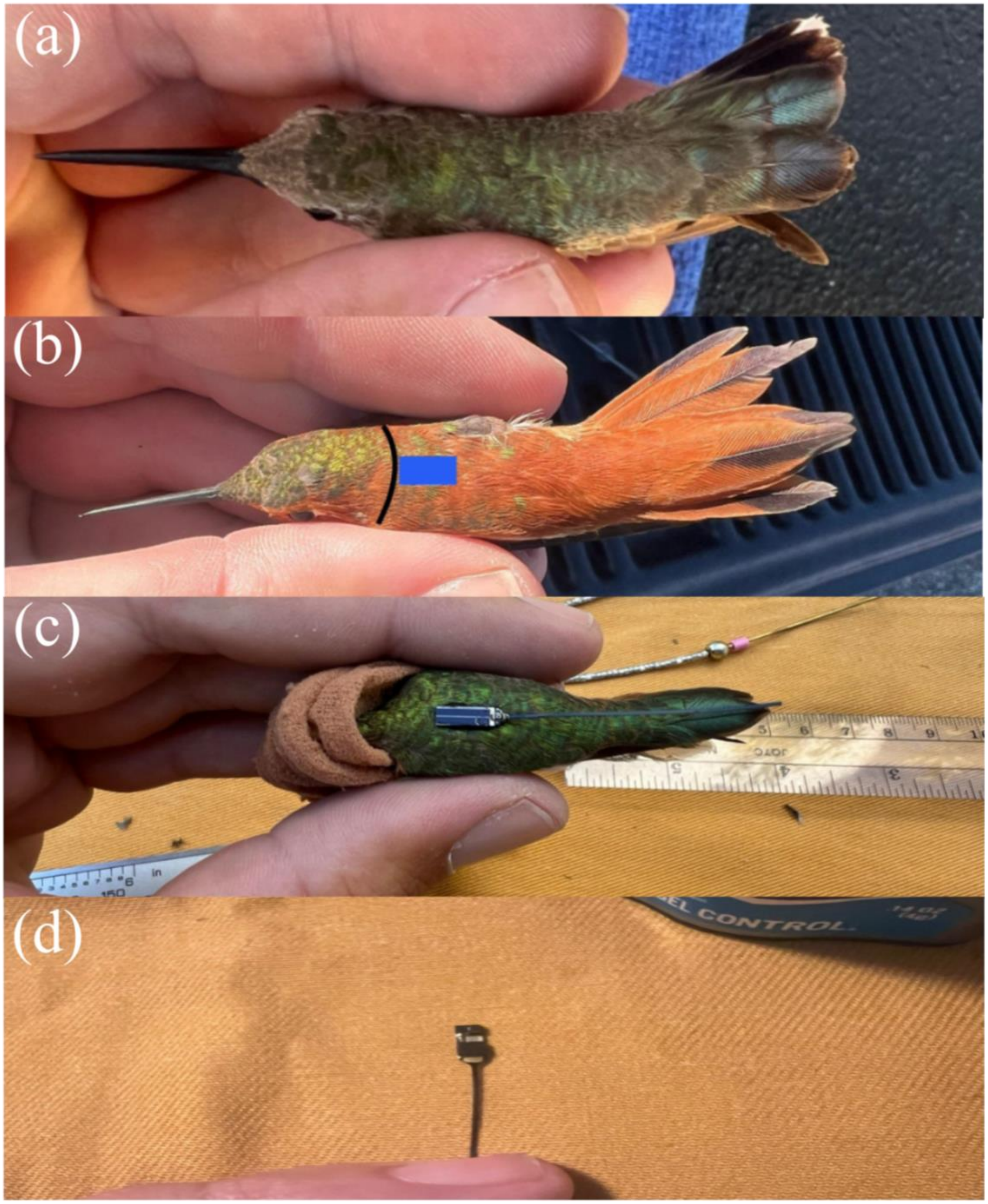
(a)
(b)
(c)
(d)

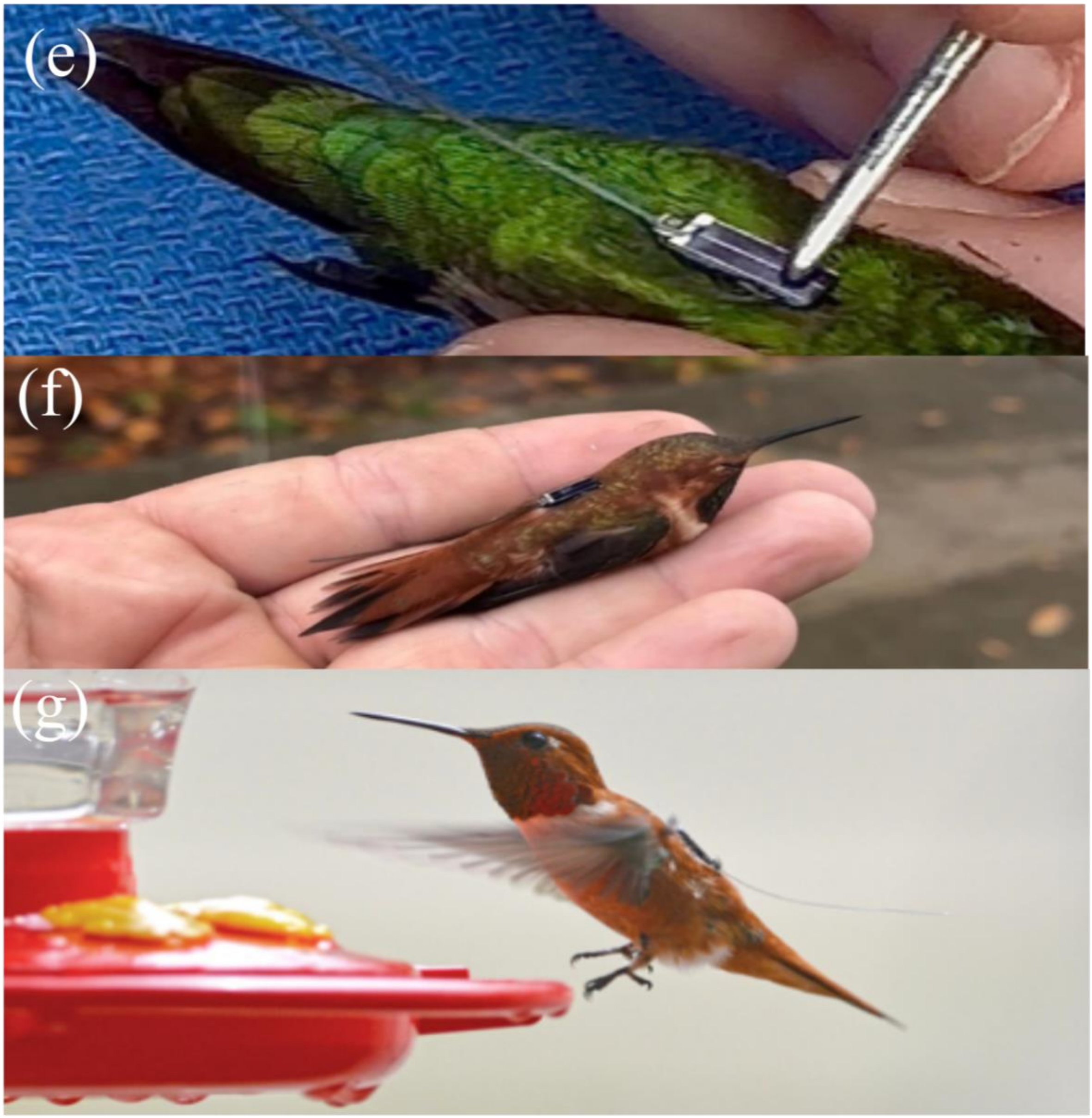
(e)
(f)
(g)

**Figure 1 (a-g).** Transmitter Application Step-by-Step Guide. The hummingbird should be held in a cigar hold with the bill facing the palm of your hand **(a).** Locate the tips of the nape feathers indicated by the black line in the photo **(b).** The tips of these feathers should mark the top of your transmitter. Using a sharp pair of cuticle scissors, trim the iridescent portions of each feather in the center of the back (interscapular region) where the transmitter will be applied, represented by the blue rectangle **(b).** We do not recommend trimming the feathers all the way to the base. The goal is to leave all feather bases intact for the glue to attach to rather than gluing directly on the skin. The hummingbird's head should be covered with a light cloth or stocking **(c).** This helps to keep the bird calm and still and helps prevent possible eye and respiratory irritation due to off-gassing super glue. Place the transmitter, solar panel down, on your table **(d).** Pin the antenna of the transmitter to the table with the pinky of the hand you are holding the bird with. Using your free hand, uncap the Gel Control super glue and apply enough to the back side of the transmitter to cover all components on the circuit board and fill the low spot in the center. Once the glue has been applied, pick up the transmitter by the antenna and set it in place on the bird with the antenna running down the center of the back toward the tail. *IMPORTANT NOTE: The oil on birds feathers rapidly cures super glue. Once it is on the bird, adherence to feathers is near instantaneous in warm weather so there are no second chances. Make sure you put it exactly where you want it.* Press the transmitter firmly to the back using a knitting needle or tapestry needle (or other smooth metal object) as sown **(e).** Important Note: DO NOT USE YOUR FINGER OR A CLOTH as the glue will stick to it and dry instantly After the transmitter is pressed firmly into place, use your knitting or tapestry needle to wipe away any squeeze out (extra glue that squeezes out the sides of the transmitter). To do this effectively and without smearing glue to other parts of the bird, the needle should be held perpendicular to the side of the body and wiped along the side of the transmitter to collect the excess glue. Wipe the excess off after addressing each side. To test if the glue has dried, you can attempt to move the end of the antenna side to side with your finger. Once the entire area around the solar panel moves as one unit, you can give the antenna a light pull. The area of the back around the transmitter should move without the transmitter pulling off. Once the glue is dry, uncover the head and release the bird (f). Again, some semblance of abnormal behavior (eyes closed and a reluctance to leave) is common when using this methodology. Encourage the bird to leave either by blowing on it from the tail of the bird, wiggling your hand to elicit a reaction, or by changing its position in your hand (f). Post-release monitoring should occur, if possible, even if it is just watching the bird fly until it is out of sight (g). **For More information and to learn more about "HumTrack", visit https://www.bandingcoalition.org/humtrack.**
